# Selective plane activation structured illumination microscopy

**DOI:** 10.1101/2023.06.19.545558

**Authors:** Kenta Temma, Ryosuke Oketani, Toshiki Kubo, Kazuki Bando, Shunsuke Maeda, Kazunori Sugiura, Tomoki Matsuda, Rainer Heintzmann, Tatsuya Kaminishi, Koki Fukuda, Maho Hamasaki, Takeharu Nagai, Katsumasa Fujita

## Abstract

Resolution enhancement in structured illumination microscopy is hindered when volumetric samples are observed owing to background signals from out-of-focus planes. In this study, we utilized selective plane illumination and reversibly photoswitchable fluorescent proteins to realize structured illumination within the focal plane and eliminate the out-of-focus background. We demonstrated high-resolution fluorescence imaging of volumetric samples, including dense fluorescent objects in a cell spheroid.

## Main

Fluorescence microscopy has expanded its imaging capability by spatially or temporally modulating the fluorescence emission. For example, the spatial modulation of illumination light, such as in laser scanning^1^, stimulated emission depletion^2^, and selective plane illumination microscopy^3^, employ Gaussian, doughnut, or sheet illumination to localize the emission area. Temporal modulation also emerges in fluorescence emissions, such as saturated excitation^4^, stepwise excitation^5–7^, and photoactivated localization^8^ microscopy, to induce or extract nonlinear fluorescence responses for spatial resolution improvement.

Structured illumination microscopy (SIM) utilizes spatial modulation and temporal pattern switching simultaneously in the illumination light for super-resolution imaging^9,10^. Spatial modulation with a sinusoidal pattern allows the detection of high-spatial-frequency information that is not accessible in conventional imaging optics. Resolution enhancement can also be achieved in the depth direction using three-dimensional (3D) modulation patterns.

However, imaging thick and dense samples remains challenging because out-of-focus signals reduce the contrast of the acquired images and cause significant artifacts in the image reconstruction process. Several combinations of SIM and selective plane illumination (SPI) have been employed to address this issue. One study utilized a structured light sheet formed with a spatial light modulator to illuminate samples from a side objective lens, yielding depth discrimination and one-directional resolution improvement^11^. The other is the addition of oblique illumination with a rotational scanner, which achieves an isotropic improvement in spatial resolution^12^. These methods effectively utilize the pupil of an illumination objective lens to achieve SI and SPI simultaneously; however, this integrated illumination limits the available numerical aperture (NA) for SI or SPI, which limits the benefit of the super-resolution technique.

In this paper, we propose selective plane activation structured illumination microscopy (SPA-SIM), wherein selection of the observation plane by sheet illumination and fluorescence excitation by structured illumination are separately carried out by two mutually orthogonal objective lenses. The separation of plane selection and excitation allowed us to arbitrarily choose illumination patterns in the light sheet and structured illumination independently without being constrained by the single-objective NA. To realize this separation, we introduced reversibly photoswitchable fluorescent proteins (RSFP). The emissive capability of the fluorescent protein is turned on by sheet-illumination light to select the observation plane, and subsequent fluorescence excitation by structured illumination realizes a structured emission pattern from a thin layer.

We confirmed the imaging properties of SPA-SIM and its advantages in the observation of thick samples by calculating the effective point spread functions (PSF) and images of fluorescent beads distributed in three dimensions. We also experimentally observed biological samples, ranging from single cells to volumetric spheroids, to confirm the improvement in image contrast and resolution in SIM by selective plane activation. In this study, we evaluated several illumination modes and found that two modes were effective for specific measurements. Single-photon activation with a light sheet provides relatively efficient photoactivation, enabling 1-frame/s 2D observation, which is sufficient for 3D observation of living cells. Two-photon activation with a scanned Bessel beam (Bessel sheet) and three-dimensional structured illumination succeeded to observe the interior of a cell spheroid with a diameter of 100 µm at the depth of up to 43 µm. Both theoretical and experimental demonstrations confirmed that the SPA-SIM is beneficial for the observation of a wide variety of specimens, including thick and dense samples, with the capability of choosing optimal illumination combinations depending on the purpose of observation.

SPA-SIM encompasses optical sectioning capability and high spatial resolution owing to the integration of SPIM and SIM via the RSFP (Fig. 1a). SPIM obtains a thin fluorescence distribution excited by plane illumination localized within the focus of the detection objective, which reduces the out-of-focus signal and improves the axial resolution. SIM utilizes multibeam interference to create a sinusoidal pattern at the focal plane and retrieves high spatial frequency information for super-resolution. We exploited the advantages of these techniques assisted by the RSFP, which can apply two different illumination patterns to determine the emission pattern in a sample. Selective plane activation localizes the on-state protein and structured illumination excites it for observation, realizing a thin, sinusoidal emission pattern inside the sample while suppressing out-of-focus excitation. Consequently, SPA-SIM provides a high 3D spatial resolution, even in a dense volumetric sample.

**Fig. 1.**
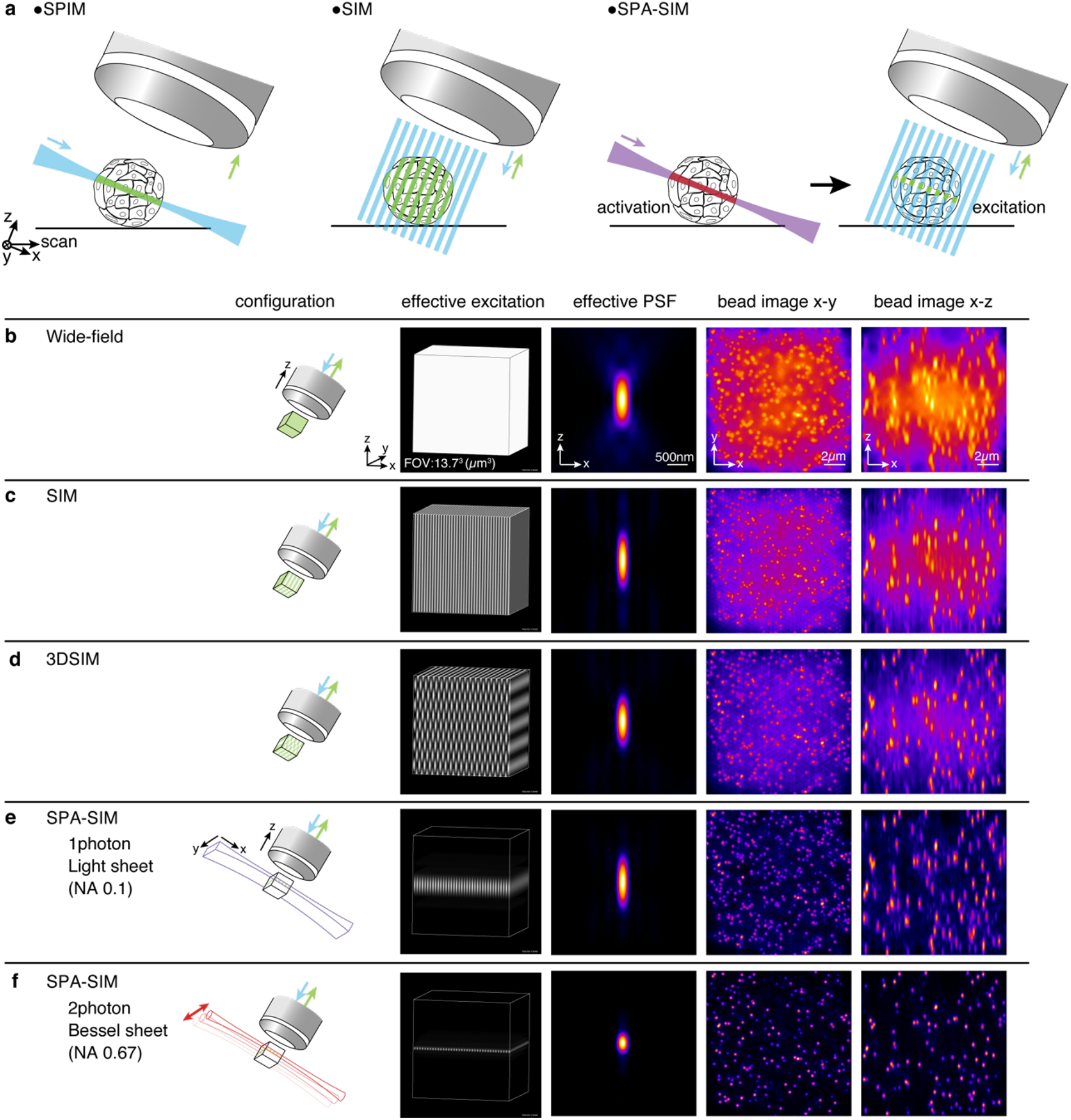
(a) Imaging schemes of selective plane illumination microscopy (SPIM), structured illumination microscopy (SIM), and selective plane activation structured illumination microscopy (SPA-SIM). Comparisons of fluorescence emission patterns, effective PSFs, simulated images of fluorescent beads scattered in a volume with (b) wide-field microscopy, (c) SIM, (d) 3DSIM, (e) SPA-SIM using single-photon light-sheet activation, and (f) SPA-SIM using two-photon scanned-Bessel-beam activation. An effective NA of 0.1 for the light sheet was chosen in (e) to set the activation area (x-y) comparable to that in the scanned Bessel beam activation (NA:0.67). A homogeneous fluorescent sample was assumed for the calculation of fluorescence emission to estimate the effective excitation patterns. The effective PSFs for (c-e) were calculated including image formation and SIM reconstruction. For the generation of three-dimensional fluorescence images, 5000 beads with a diameter of 200 nm distributed randomly in a volume of 13.7×13.7×13.7 µm^3^ were assumed to compare the spatial resolution and the capability of suppressing out-of-focus signals.

To verify our concept, we numerically calculated the imaging properties of the SPA-SIM with different modes of selective plane activation. We calculated effective excitation patterns and effective PSFs, and simulated the imaging of 5000 fluorescent beads with a diameter of 200 nm three-dimensionally distributed in a volume of 13.7×13.7×13.7 µm^3^. Fig. 1b and 1c confirm that SIM provides an improvement in the lateral spatial resolution and a slight improvement in the axial resolution compared to wide-field microscopy. However, in volumetric sample images, out-of-focus signals degrade the image contrast and induce significant artifacts during the image reconstruction process. The suppression of out-of-focus signals with the 3DSIM (Fig. 1d) was still insufficient to remove all imaging artifacts. In contrast, selective plane activation drastically improved image contrast and reduced artifacts (Fig. 1e-f). Although the effective PSF using single-photon activation does not show a strong improvement in axial resolution, owing to the low effective NA (0.1) of the illuminating light sheet to warrant a sufficiently large field of view (FOV), the simulated fluorescence image confirmed a drastic improvement in image contrast and artifact removal, presumably owing to the suppression of out-of-focus signals. Two-photon activation with a scanned Bessel beam provides a higher spatial resolution, background rejection capability, and a sufficiently large FOV owing to its nonlinear activation capability and full use of the sheet illumination NA (0.67). The effective PSF in Fig. 1f shows an isotropic spatial resolution, which was also confirmed in the imaging simulation of fluorescent beads. We also confirmed that selective plane activation improves the contrast of the structured illumination in a dense sample, as shown in Supplementary Fig. 2, which reduces artifacts in the reconstructed SIM images owing to high background signals. Detailed comparisons of PSFs and OTFs are presented in Supplementary Fig. 3, which clearly shows the improvement in the axial resolution in SPA-SIM.

**Fig. 2.**
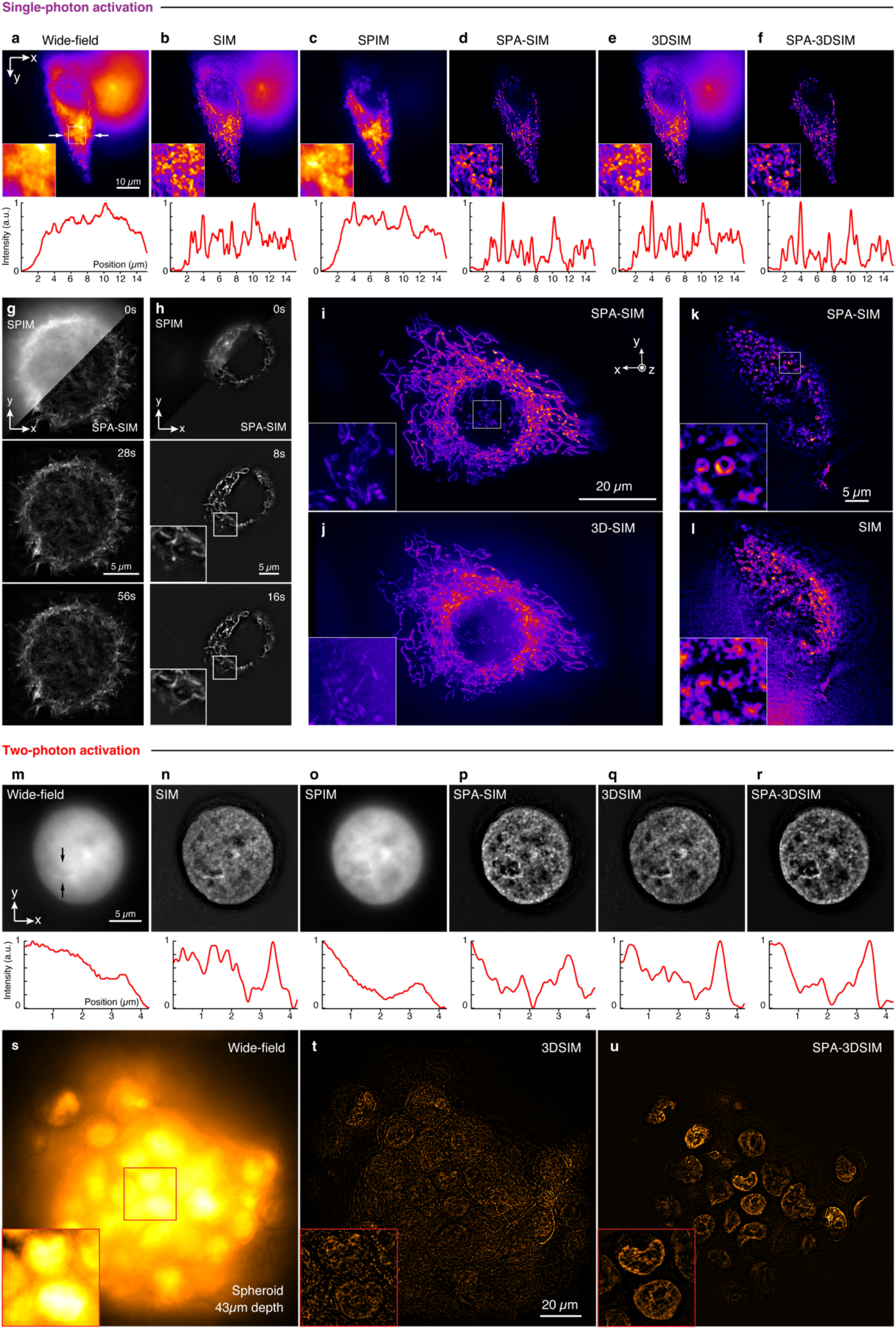
(a-f) Fluorescence images of mitochondria in fixed HeLa cells and intensity profiles between white arrowheads. The images were taken and reconstructed in the imaging modes of (a) wide-field, (b) SIM, (c) SPIM, (d) SPA-SIM, (e) 3DSIM, and (f) SPA-3DSIM. Time-lapse images of (g) actin filaments and (h) mitochondria of a living HeLa cell with SPIM (first frame) and SPA-SIM modes. 3D-rendered images of mitochondria in a living HeLa cell with (i) SPA-SIM and (j) 3DSIM. Fluorescence images of autophagosomes in living mouse embryonic fibroblast (MEF) cells obtained in (k) SPA-SIM and (l) SIM modes. In (a) - (l), the samples were labeled by Skylan-NS and activated by the 405 nm with light-sheet (c, d, f, g, h, i, k) and wide-field (a, b, e, j, l) illumination. (m-r) Fluorescence images of histone H2B in a nucleus of a living HeLa cell labeled by rsGamillus-S and the intensity profiles between black arrowheads. (m-r) are obtained under the same imaging modes as (a-f), but a scanned Bessel beam with a wavelength of 780 nm was applied for two-photon activation in (o, p, r). Nuclei in a cell spheroid labeled by rsGamillus-S with a diameter of ∼100 µm were observed with (s) wide-field, (t) SIM, and (u) SPA-3DSIM at the depth of 43 µm. The details of the imaging conditions are described in Extended data Fig. 2.

In our study, we examined different combinations of illumination patterns for SPA-SIM, as shown in Extended Data Fig. 1, and concluded that both single-photon activation with low-NA sheet illumination and two-photon activation with a scanned Bessel beam are suitable for observing volumetric specimens with respect to their uniformity in the activation pattern and background rejection capability. These activation patterns maintained their uniformity, at least within the FOV of the SIM. We also summarize the imaging properties of 3DSIM with selective plane activation in Supplementary Fig. 3. The 3DSIM further enhanced the axial discrimination capability of the SPA-SIM, with a slight decrease in the lateral resolution.

As the first demonstration of our concept, we used a self-made SPA-SIM setup (Supplementary Fig. 4) and observed mitochondria labeled with Skylan-NS^13^ that exhibited a high switching contrast in fixed HeLa cells in six different imaging modes to compare the image contrast and spatial resolution (Fig. 2a-f). In the wide-field mode (Fig. 2a), dense mitochondrial distributions were not visible because of low spatial resolution and high background signals. SIM (Fig. 2b) improved the spatial resolution, but out-of-focus signals degraded the contrast of the image and visualization of intracellular structures. The selective-plane illumination mode (Fig. 2c) reduced the out-of-focus signals, whereas the lateral spatial resolution did not improve. By contrast, the SPA-SIM (Fig. 2d) mode combines the advantages of both selective plane illumination and SIM, exhibiting drastic improvements in image contrast and spatial resolution, even for a densely labeled thick specimen. A comparison with the results of 3DSIM^14^ (Fig. 2e) confirmed that SPA-SIM reduced the influence of the out-of-focus background more effectively, which benefits the visualization of intracellular structures. We also applied selective plane activation to the 3DSIM and confirmed further improvements in its optical sectioning capability (Fig. 2f). The out-of-focus light in the image by the SPA-3DSIM was reduced more efficiently than that by the SPA-SIM, while the practical spatial resolution was slightly inferior to that of the SPA-SIM, as can be seen in the PSFs of the SIM and 3DSIM in Fig. 1 and Supplementary Fig. 3.

Fig. 2g-h show the results of time-lapse imaging of actin (Supplementary Movie1) and mitochondria (Supplementary Movie2) in HeLa cells at 0.6 and 1 frames/s with an interval of 7 and 4 s, respectively. SPA-SIM visualizes actin movement and mitochondrial fusion at a higher spatial resolution than SPIM. We also observed the three-dimensional distribution of mitochondria in living HeLa cells with SPA-SIM and 3DSIM, as shown in Fig. 2i-j. SPA-SIM provided higher contrast and better visualization of the mitochondria distributed throughout the cell, including those obscured by the background in 3DSIM. A comparison between the SPA-SIM mode and SIM for three-dimensional imaging is shown in Supplementary Fig. S6.

We also observed autophagosomes labeled with the autophagy marker protein LC3B fused to Skylan-NS^12^ in living mouse embryonic fibroblasts (MEF) using SPA-SIM and SIM, as shown in Fig. 2k-l. Typically, MEFs have a volume larger than that of HeLa cells, and thus, higher background signals result from fluorescent proteins. Owing to the rejection of out-of-focus signals, SPA-SIM successfully visualized ring-shaped structures attributable to autophagosomes that emerged upon starvation induction, whereas conventional SIM and wide-field mode failed (Supplementary Fig. 7).

For the observation of larger samples using SPA-SIM, two-photon activation using a scanned Bessel beam is suitable because it realizes a wider FOV and a higher axial resolution, as discussed in a report on Bessel beam plane illumination microscopy^15^. We applied two-photon activation using a scanned Bessel beam to selective plane-structured illumination to observe histoneH2B in living HeLa cells labeled with rsGamillus-S. HistoneH2B densely distribute in the cell nuclei. We chose rsGamillus-S because of its exceptionally high on-switching rate among the RSFPs^16^. As in the single-photon activation experiment, six observation modes were compared (Fig. 2m-r). In the wide-field mode (Fig. 2m), the structures inside the nucleus were not visible because of the high background signal. Several structures in the nucleus were observed in SPIM mode (Fig. 2o), but the lateral and axial resolutions were insufficient. Conventional SIM (Fig. 2n) and 3DSIM (Fig. 2q) improved the spatial resolution and optical sectioning capability; however, internal structures were still buried in the background signals. SPA-SIM using two-photon activation with a Bessel sheet enabled the observation of the internal structures of the nucleus with high spatial resolution and image contrast (Fig. 2p, r). In particular, the groove-like structure between the black arrowheads is best visualized in high contrast with the SPA modes.

We also observed cell spheroids to evaluate the performance of the SPA-SIM by observing a thicker sample (Fig. 2s-u). A living HeLa-cell spheroid of about 100 µm diameter with histoneH2B labeled by rsGamillus-S was observed. The observation depth was about 43 µm from the spheroid surface. In general, observing cells in a spheroid using SIM is difficult because the out-of-focus fluorescence obstructs the detection of the in-focus fluorescence and reduces the contrast of the structured patterns. We experimentally confirmed that the wide-field and 3DSIM modes were difficult to use for imaging the internal structure of the spheroid, as shown in Fig. 2s-t. In contrast, SPA-3DSIM achieved super-resolution imaging of histoneH2B distribution in a spheroid (Fig. 2u). We also compared SPIM and SPA-3DSIM, as shown in Supplementary Fig. 8, to evaluate resolution improvement.

To discuss our findings, we first compared SPA-SIM with conventional 3D-SIM. SPA-SIM further reduces the out-of-focus signal and achieves a greater imaging depth for dense objects. SPA-SIM also requires fewer raw images for a single 3D super-resolution image; however, each raw image requires longer irradiation, including the activation and deactivation of the RSFPs, which affects the imaging speed. Although longer irradiation times can also be harmful to the samples. However, we did not observe significant cell damage using SPA-SIM imaging. In cell imaging, irradiation at wavelengths below 450 nm is more harmful than that at longer wavelengths. The use of light sheet illumination for activation possibly reduced cell damage.

The choice of RSFPs is important for achieving SPA-SIM. In the single-photon activation mode, RSFPs with a high switching rate and switching contrast are suitable for observations with high temporal resolution, such as Skylan-NS, Dronpa, rsGamillus-F, and other RSFPs with similar properties. In two-photon activation mode, rsGamillus-S was superior. Although two-photon excitation is less efficient than single-photon excitation and leads to low switching efficiency, the high on-switching rate of rsGamillus-S provides sufficient signals within short activation periods, as confirmed by comparison with Skylan-NS in Supplementary Fig. 10. In this study, we used only negatively switchable RSFPs; however, positive types were also available using different wavelengths of light for SPA and SIM. Protein engineering plays an important role in rapid imaging. The development of fluorescent proteins with optimized quantum yield, switching rate, and absorption cross-section boosts the imaging speed.

Sample-induced aberrations are a problem in deep imaging. In our SPA-SIM system, the aberrations make imaging deeper parts difficult and limit the imaging depth to about 43µm in a spheroid of ∼100 µm diameter (Fig. 2u). Computational approaches such as Blind-SIM^17^ should be able to deal with aberrations in SIM illumination in SPA-SIM. The implementation of adaptive optics in SPA-SIM should also be useful for maintaining both the selective plane and structured illumination, correcting the aberrations on the detection side, and accessing deeper region ^18,19^.

The axial resolution of the SPA-SIM can be further improved by the arbitrariness of the illumination. Pupil engineering expands the useful choices of illumination patterns for sheet activation. A sectioned Bessel beam^20^ can suppress the side lobes in Bessel beam illumination. We expect a high spatiotemporal resolution with single-photon-sectioned Bessel beam activation. A lattice light sheet^11^ is also a suitable candidate for SPA-SIM to further improve the axial resolution. Using visible wavelength for two-photon activation also improves the axial resolution with a thin sheet ^21,22^.

The SPA-SIM principle can be applied to single-objective systems. Although the axial resolution is lower than that of two-objective systems, it provides background reduction that is beneficial for SIM with easier sample handling and implementation.

## Methods

### Optical setup

Our imaging system consists of a sheet illumination component for activation and a structured illumination component for excitation. For the sheet illumination part, we form a light sheet or a Bessel light sheet, which is realized using a cylindrical lens or an axicon lens with a scanning mirror, respectively. The light sheet is relayed to the sample space by relay lenses and a water immersion objective lens (28.6×, NA0.67 model 54-10-7, Special Optics). As a structured illumination part, a 2D or 3D stripe pattern is produced by a spatial light modulator (SLM, SXGA 3DM, Fourth dimension display) for super-resolution imaging. The patterned light is relayed and overlapped with the sheet illumination using another water immersion objective lens (25×, NA1.1, CFI75, Nikon) arranged orthogonally to the objective lens for sheet activation. The fluorescence emission is collected using the same objective as the structured illumination and imaged using an EMCCD camera (OLCA-Flash 4.0 v3, Hamamatsu Photonics). The details of the setup are shown in Supplementary Fig. 4.

### Reconstruction

The reconstruction of super-resolution images for SPA-SIM is essentially the same as that for conventional SIM (see the mathematical descriptions in the supplementary information). Pre-processing for acquired raw images and SIM reconstruction are performed with a self-made program in MATLAB, and partially with ‘fairSIM’^23^, a plugin of ImageJ software (Fig. 2i-j). For preprocessing, normalization of the fluorescence intensity among images and deskewing for the 3D image stack were applied to the series of raw images. A small sample drift between frames was compensated through phase correlation. In SIM reconstruction, a spatial Fourier transform and matrix-based unmixing are performed on the preprocessed images to separate and reassign the high spatial frequency information in the frequency space. We optimized the reconstruction parameters, such as the phases and frequencies of structured illumination, using preprocessed raw images to avoid reconstruction artifacts owing to the mismatch between the experimental results and the theoretical estimation. The optimization was based on the cross-correlation of the frequency components explained in ref ^24^. We applied a Wiener filter and weighted averaging^25^ to each frequency component to mitigate Poisson noise in the frequency space and enhance the high-frequency information carried by the structured illumination. The resultant frequency distribution was inverse Fourier-transformed to obtain a reconstructed super-resolution image.

### Sample Preparation

#### Plasmid constructs

The pcDNA3-Lifeact-Skylan-NS plasmid was used as described in a previous study^14^. The mitochondria-localized Skylan-NS construct was created by replacing the rsGamillus sequence of pcDNA3-Cox-VIII ×2 -rsGamillus used in our previous study^14^ with the Skylan-NS sequence. The construct for the nuclear localization of rsGamillus-S was created by replacing the Gamillus sequence of pcDNA3-Gamillus-H2B created in a previous study^26^ with the rsGamillus-S sequence. Sequence substitution was performed using the hot fusion method^27^.

### Cell culture and transfection

HeLa cells were cultured in high-glucose Dulbecco’s modified Eagle medium (Wako, DMEM, 043-30085) supplemented with 10 % (v/v) fetal bovine serum (Biowest, S1400-500). To express fluorescent proteins, plasmid of (Skylan-NS and rsGamillus-S) were injected into the cell suspension mixed with Opti-MEM and electroporated using an electroporator (Nepagene, NEPA21) according to the manufacturer’s instructions. After transfection, HeLa cells were seeded onto glass substrates.

### Fixed cell specimens for Fig. 2a-f

HeLa cells were fixed after transfection with fluorescent proteins and seeded onto glass substrates. Cells were first washed three times with PBS to remove DMEM, treated with 4 % paraformaldehyde in PBS for 15 min, and rinsed in PBS three times again.

### Autophagosome in MEF for Fig.2k-l

The coding sequences of Skylan-NS^12^ (Addgene, plasmid #86785) and LC3B were subcloned into the pMRX-IRES-puro vector and transduced into MEFs using retroviruses packaged in Plat-E cells, as previously described^28^. MEFs stably expressing Skylan-NS–LC3B were selected with 3 µg/mL puromycin (Invivogen), seeded on a glass slide coated with 0.3 mg/ml collagen (Nitta Gelatin, Cell matrix Type I-C), and grown overnight at 37 °C with 5 % CO_2_ in high-glucose Dulbecco’s modified Eagle medium (Sigma, D6429) supplemented with 10 % (v/v) heat-inactivated fetal bovine serum (Sigma, F7524) and 2 mM L-glutamine (Sigma, G7513). The medium was then replaced with Earle’s Balanced Salt Solution (Sigma, E3024) with 125 nM Bafilomycin A1 (Sigma, B1793) to induce starvation and inhibit autophagosome-lysosome fusion, respectively. After 2 h of incubation, cells were placed in PBS and observed.

### Cell spheroid in Fig. 2s-u

Cell spheroids consist of HeLa cells with rsGamillus-S-H2B stably expressed on their H2B, which was picked up from the transfected HeLa cells by culturing in a mixture of 1 % geneticin and DMEM on a 90 mm dish to eliminate the cells without expressing fluorescent proteins. Spheroids were formed using a 35-mm dish with a low-adhesive micro-patterned bottom (Iwaki, Ezsphere, 4000-900). HeLa cells were seeded in dishes at a density of 50 cells/spheroid. Cells were incubated at 37 °C in an atmosphere of 5 % CO_2_ for 2-3 days to form spheroids. When the diameter of the spheroids reaches around 100 µm, they were collected and placed on glass substrates. After 2 h of adhesion to the substrates, the spheroids were observed using SPA-SIM.

## Data availability

The data underlying the results presented in this paper are not publicly available at this time but may be obtained from the authors upon reasonable request.

## Code availability

The MATLAB-based SIM reconstruction code is available from the authors upon request.

## Acknowledgements

The authors thank Shogo Kawano (Osaka University) for the help in developing the control system for the SPA-SIM setup. The authors thank Dr. Tamotsu Yoshimori (Osaka University) for providing facilities and cells necessary for the establishment of the Skylan-NS–LC3B expressing MEF. The authors thank Dr. Shoji Yamaoka (Tokyo Medical and Dental University) and Dr. Toshio Kitamura (The University of Tokyo) for providing the pMRX-IRES-puro retroviral vector and Plat-E retroviral packaging cells, respectively. This work was partially supported by Core Research for Evolutionary Science and Technology by Japan Science and Technology Agency (grant no. JPMJCR15N3 to T.N. and JPMJCR1925 to K.F.). This work was also partially supported by grant-in-aid from the Ministry of Education, Culture, Sports, Science and Technology, Japan (grant no. 18H05410 to T.N.). A part of this work was performed under the Research Program of “Dynamic Alliance for Open Innovation Bridging Human, Environment and Materials” in “Network Joint Research Center for Materials and Devices” to K.F. and T.N.

## Author contributions

K.T. and R.O. contributed equally to this study. K. Fujita supervised the project and established the SPA-SIM concept. R.O. built the numerical calculation programs and K.T. modified the program and performed the calculations. R. O., K. T., K. B., T. Kubo, and S. M. designed the optical setup. R. O. and K. T. constructed the setup. T. Kubo partially developed the control system. K. T. and R. O. performed the experiments. K.T. and T. Kubo prepared the single-cell and cell spheroid samples. K.S., T.M., and T.N. prepared reversibly photoswitchable fluorescent proteins for SPA-SIM measurements. K. Fukuda, T. Kaminishi, and M.H. prepared MEF for autophagosome imaging. R.H. built the reconstruction program. K.T., R.O., and T. Kubo performed reconstruction. K. T., R. O., and K. Fujita wrote the manuscript. All the authors reviewed the manuscript.

## Ethics declarations

### Competing interests

K. T., R. O., and K. Fujita filed a patent application (May 26, 2023) for the proposed method.

**Extended Data Fig. 1.**
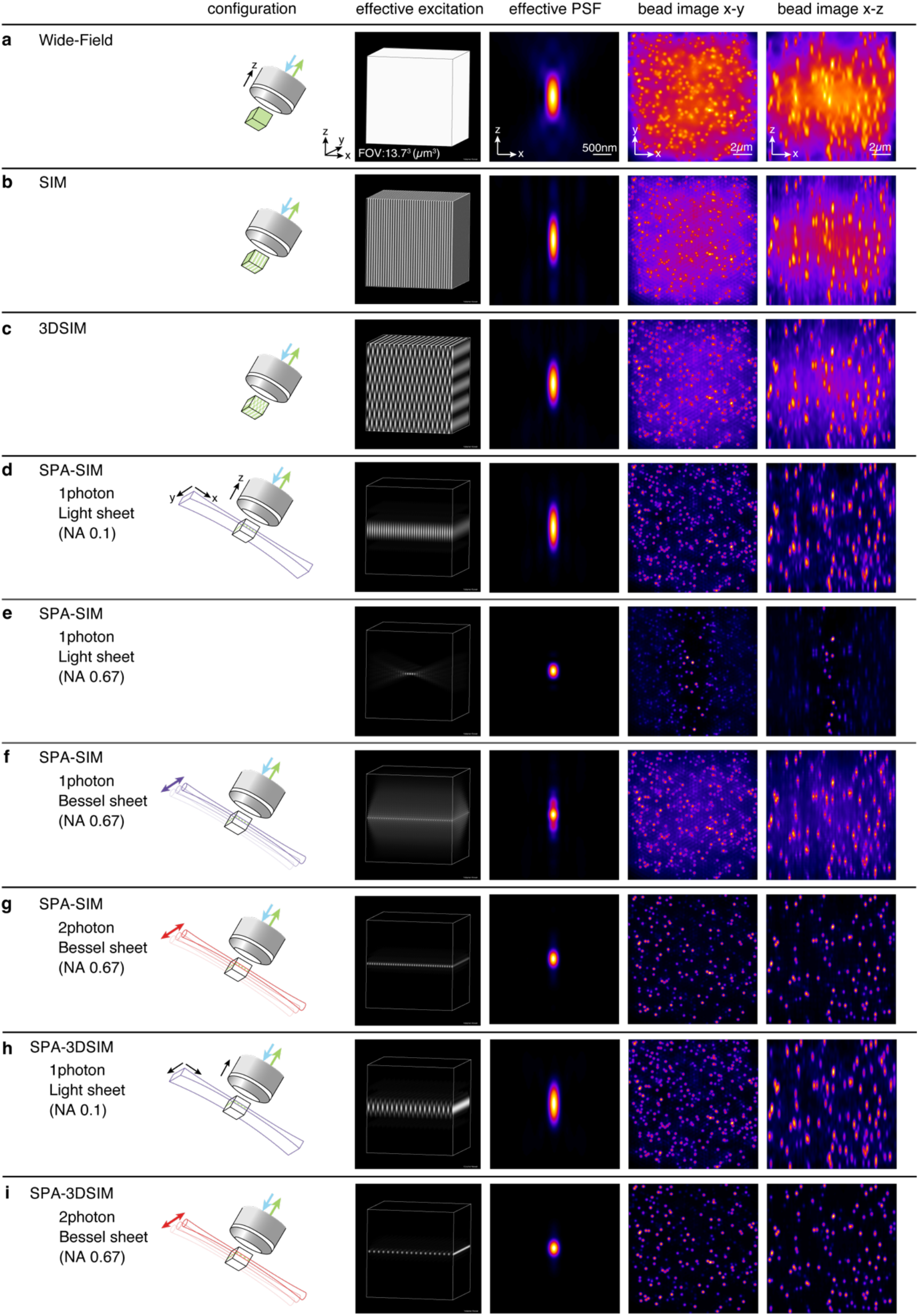
Comparison of selective plane activations. A variety of illuminations and corresponding fluorescence emissions, effective PSFs and simulated bead images in SPA-SIM. For comparison, Fig.1(b), (c), (d), (e), and (f) are shown in (a), (b), (c), (d), and (g), respectively. In addition, the cases of a light sheet with high NA and single-photon Bessel sheet for activation and SPA-3DSIM are shown in (e), (f), (h), and (i). FWHM of PSFs in axial direction are 787.0, 786.8, 726.5, 745.2, 251.4, 544.6, 300.5, 695.2, and 284.3 nm for wide-field, SIM, 3DSIM, SPA-SIM with low NA light sheet, high NA light sheet, single-photon activation Bessel sheet, and two-photon activation Bessel sheet, SPA-3DSIM with low NA light sheet, and two-photon activation Bessel sheet, respectively. In lateral direction, FWHM of PSFs are 247.5, 168.7, and 184.5 nm in conventional wide-field, SIM and 3DSIM. NA of 1.1 was assumed for excitation and detection. Structured illumination was achieved using 60 % of full NA during excitation.

**Extended Data Fig. 2.**
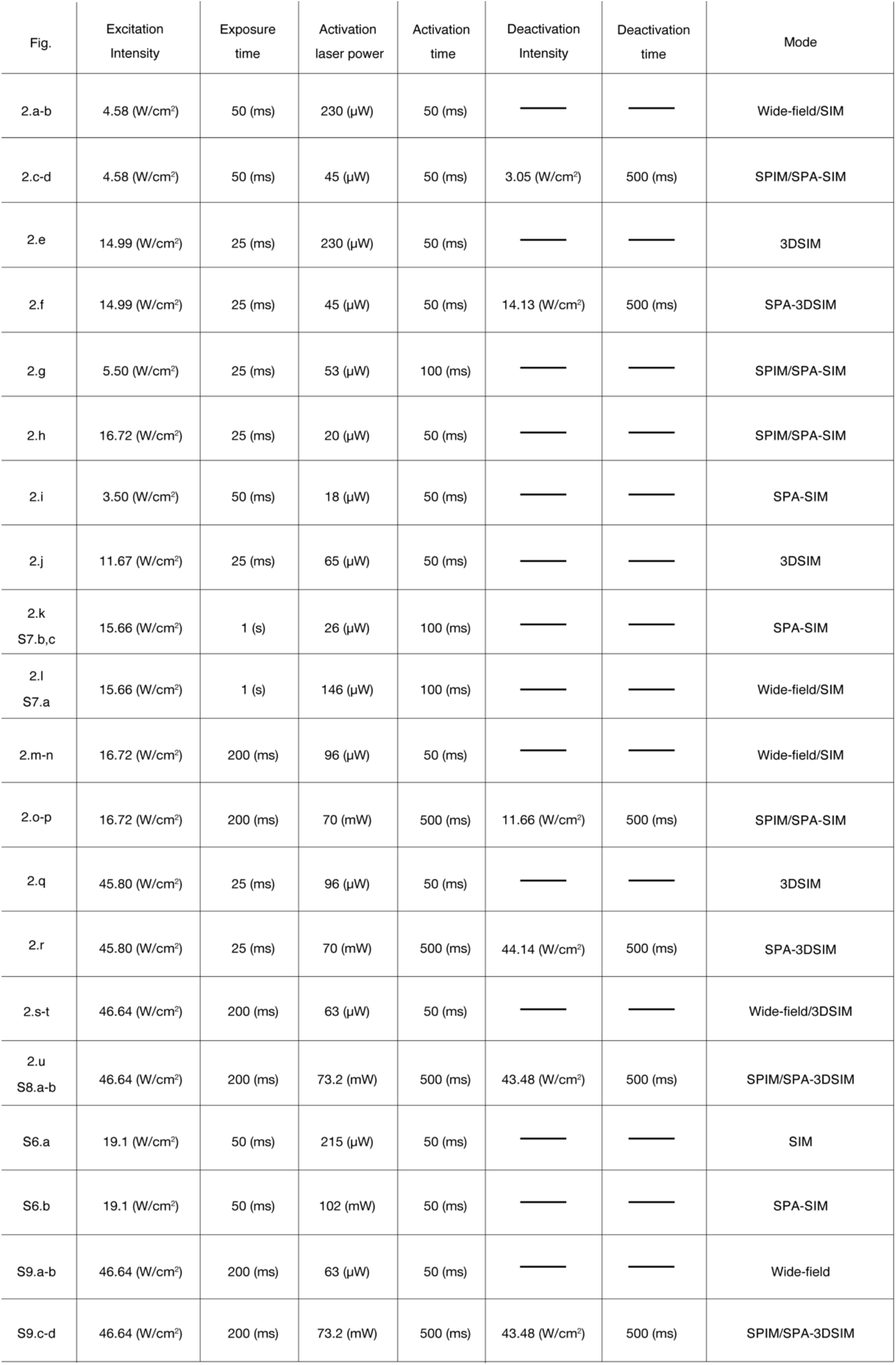
Imaging conditions for acquired images in Fig. 2. and Supplementary Figures.

