## Supplementary Information for "Selective plane activation structured illumination microscopy"

### S.1 Theoretical calculation

#### Light distribution for SPA-SIM

The electric field distribution $E(\boldsymbol{r})$ of light after passing through a lens is represented as an integral of plane waves as follows:

$$E(\boldsymbol{r}) = \int_{-\infty}^{\infty} P(\boldsymbol{r}')\mathrm{e}^{\mathrm{i}(\boldsymbol{k}(\boldsymbol{r}')\cdot\boldsymbol{r}-\omega t)}\mathrm{d}\boldsymbol{r}', \qquad (1)$$

where $\boldsymbol{r}$ and $\boldsymbol{r}'$ are the position vectors in the object space and pupil plane, respectively; $P(\boldsymbol{r})$ is the complex-valued pupil transmission function of the lens; $t$ is the time; and $\boldsymbol{k}(\boldsymbol{r})$ and $\omega$ are the wave number vector and angular frequency, respectively. When illumination for activation is provided by a lens and objects including photoswitchable fluorophores are placed near the focus of the lens, the distribution of the activated fluorophores is represented as

$$c_{\mathrm{act}}(\boldsymbol{r}) = \left|E_{\mathrm{act}}(\boldsymbol{r})\right|^2 o(\boldsymbol{r}), \qquad (2)$$

where $E_{\mathrm{act}}(\boldsymbol{r})$ and $o(\boldsymbol{r})$ are the illuminating electric field distribution for the activation and distribution of the object, respectively. The activated fluorophores were then excited by a second illumination. The resultant distribution of the excited fluorophores is described as

$$c_{\mathrm{ex}}^{nm}(\boldsymbol{r}) = \left|E_{\mathrm{ex}}^{nm}(\boldsymbol{r})\right|^2 c_{\mathrm{act}}(\boldsymbol{r}), \qquad (3)$$

where $E_{\mathrm{ex}}^{nm}(\boldsymbol{r})$ is the electric field distribution of the structured excitation illumination, and $n, m = 1\cdots 3$ are the indices of the angle and phase of the illumination pattern for the SIM reconstruction. For two-dimensional (2D) SIM, the intensity distribution of the structured illumination is represented as

$$\left|E_{\rm ex}^{nm}(\boldsymbol{r})\right|^2 = 1 + \frac{1}{2}\mathrm{e}^{\mathrm{i}(\boldsymbol{k}_n\cdot\boldsymbol{r}+\phi_{nm})} + \frac{1}{2}\mathrm{e}^{-\mathrm{i}(\boldsymbol{k}_n\cdot\boldsymbol{r}+\phi_{nm})}, \tag{4}$$

where $\boldsymbol{k}_n$ is the wave vector, and $\phi_{nm}$ is the phase of a sinusoidal illumination pattern with an angle-dependent global phase. Fluorescence emitted from the excited fluorophores was measured using detection optics. The distribution of the detected fluorescence $I(\boldsymbol{r})$ was

$$I^{nm}(\boldsymbol{r}) = h_{\rm det}(\boldsymbol{r}) \otimes \left[\left|E_{\rm ex}^{nm}(\boldsymbol{r})\right|^2 c_{\rm act}(\boldsymbol{r})\right], \tag{5}$$

where $h_{\rm det}(\boldsymbol{r})$ is the point spread function (PSF) of the detection optics. By substituting (2) into (5), the relationship between the illumination patterns and the resultant image can be described as

$$I^{nm}(\boldsymbol{r}) = h_{\rm det}(\boldsymbol{r}) \otimes \left[\left|E_{\rm ex}^{nm}(\boldsymbol{r})\right|^2 \left|E_{\rm act}(\boldsymbol{r})\right|^2 o(\boldsymbol{r})\right]. \tag{6}$$

(6) indicates that the integration of sheet illumination for activation and structured illumination for excitation determines the distribution of excited fluorophores.

**Image reconstruction of SIM for super-resolution**

Conventional SIM reconstruction methods[1] are directly applicable to SPA-SIM, because the mechanism of image formation is the same as that of SIM. We now briefly explain the reconstruction procedure in the case of the 2DSIM.

SIM Reconstruction was performed based on the Fourier transform and matrix-based unmixing. The Fourier transform of (5) is represented as

$$\begin{aligned}\tilde{I}^{nm}(\boldsymbol{k}) &= \tilde{h}_{\rm det}(\boldsymbol{k})\left[\left|\tilde{E}_{\rm ex}^{nm}(\boldsymbol{k})\right|^2 \otimes \tilde{c}_{\rm act}(\boldsymbol{k})\right] \\ &= \tilde{h}_{\rm det}(\boldsymbol{k})\left[\tilde{c}_{\rm act}(\boldsymbol{k}) + \frac{\mathrm{e}^{-\mathrm{i}\phi_{nm}}}{2}\tilde{c}_{\rm act}(\boldsymbol{k}+\boldsymbol{k}_n) + \frac{\mathrm{e}^{\mathrm{i}\phi_{nm}}}{2}\tilde{c}_{\rm act}(\boldsymbol{k}-\boldsymbol{k}_n)\right]\end{aligned} \tag{7}$$

where $\tilde{X}$ represents Fourier transform of $X$. There are 3 unknowns: $\tilde{c}'_{\mathrm{act}}(\boldsymbol{k})$, $\tilde{c}'_{\mathrm{act}}(\boldsymbol{k}+\boldsymbol{k}_n)$ and $\tilde{c}'_{\mathrm{act}}(\boldsymbol{k}-\boldsymbol{k}_n)$ to be estimated for the reconstruction. We used 3 different phases of structured illumination in the imaging to estimate these unknowns. Then, we got,

$$\begin{bmatrix} \tilde{I}^{n1}(\boldsymbol{k}) \\ \tilde{I}^{n2}(\boldsymbol{k}) \\ \tilde{I}^{n3}(\boldsymbol{k}) \end{bmatrix} = \begin{bmatrix} 1 & \frac{1}{2}\mathrm{e}^{-\mathrm{i}\phi_{n1}} & \frac{1}{2}\mathrm{e}^{\mathrm{i}\phi_{n1}} \\ 1 & \frac{1}{2}\mathrm{e}^{-\mathrm{i}\phi_{n2}} & \frac{1}{2}\mathrm{e}^{\mathrm{i}\phi_{n2}} \\ 1 & \frac{1}{2}\mathrm{e}^{-\mathrm{i}\phi_{n3}} & \frac{1}{2}\mathrm{e}^{\mathrm{i}\phi_{n3}} \end{bmatrix} \begin{bmatrix} \tilde{h}_{\mathrm{det}}(\boldsymbol{k}) & 0 & 0 \\ 0 & \tilde{h}_{\mathrm{det}}(\boldsymbol{k}) & 0 \\ 0 & 0 & \tilde{h}_{\mathrm{det}}(\boldsymbol{k}) \end{bmatrix} \begin{bmatrix} \tilde{c}_{\mathrm{act}}(\boldsymbol{k}) \\ \tilde{c}_{\mathrm{act}}(\boldsymbol{k}+\boldsymbol{k}_n) \\ \tilde{c}_{\mathrm{act}}(\boldsymbol{k}-\boldsymbol{k}_n) \end{bmatrix} \quad (8)$$

We estimated three components for reconstruction, $\tilde{c}'_{\mathrm{act}}(\boldsymbol{k})$, $\tilde{c}'_{\mathrm{act}}(\boldsymbol{k}+\boldsymbol{k}_n)$, and $\tilde{c}'_{\mathrm{act}}(\boldsymbol{k}-\boldsymbol{k}_n)$, by solving this equation with the estimated or measured detection PSF: $\tilde{h}'_{\mathrm{det}}(\boldsymbol{k})$. $\tilde{c}'_{\mathrm{act}}(\boldsymbol{k}+\boldsymbol{k}_n)$ and $\tilde{c}'_{\mathrm{act}}(\boldsymbol{k}-\boldsymbol{k}_n)$ are shifted back to superpose the DC components in the frequency domain and are represented as $\tilde{c}'_{\mathrm{act-shifted}}(\boldsymbol{k}+\boldsymbol{k}_n)$ and $\tilde{c}'_{\mathrm{act-shifted}}(\boldsymbol{k}-\boldsymbol{k}_n)$, respectively. The estimation is performed in three different $\boldsymbol{k}_n$ for a homogenous optical passband, and the summation of these nine components is the Fourier transform of the reconstructed super-resolution image:

$$\tilde{I}_{\mathrm{SPA-SIM}}(\boldsymbol{k}) = \sum_{n=1}^{3} \tilde{c}'_{\mathrm{act}}(\boldsymbol{k}) + \tilde{c}'_{\mathrm{act-shifted}}(\boldsymbol{k}+\boldsymbol{k}_n) + \tilde{c}'_{\mathrm{act-shifted}}(\boldsymbol{k}-\boldsymbol{k}_n). \quad (9)$$

A similar reconstruction was performed using 3DSIM[2]. In particular, in 3DSIM, selective plane activation can modify the effective illumination pattern depending on the sheet thickness. This effect changes the weights of the frequency components derived from the structured illumination, which should be considered when choosing the coefficients in the unmixing process. In our reconstruction, the coefficients were optimized according to the weights of the frequency components in the raw dataset.

| | Activation | | Excitation & Detection |
|---|---|---|---|
| objective | NA 0.1<br>water (n = 1.33) | NA 0.67<br>water (n = 1.33) | NA 1.1<br>water (n = 1.33) |
| wavelength | Light-sheet<br>405nm | Light-sheet & single-photon Bessel:<br>405nm<br>two-photon Bessel:<br>780nm | Excitation:<br>488nm<br>Fluorescence:<br>522nm |

Fig. S1 Calculation conditions used in Fig. S2 and S3. (n: refractive index)

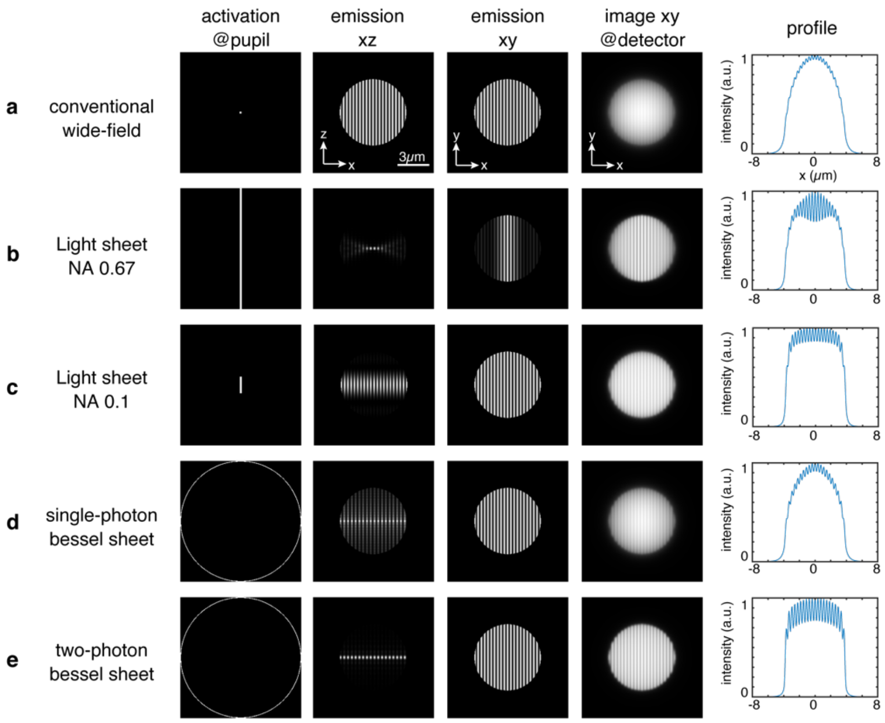


Fig. S2 Intensity distributions of activation light at the pupil of the objective lens, fluorescence emission on the xz and xy planes at the object, and the fluorescence signal on the image sensor calculated for the five imaging modes of (a) conventional SIM, light-sheet with single-photon-activation using NAs of (b) 0.67 and (c) 0.1, and Bessel sheet with (d) single- and (e) two-photon-activation using an NA of 0.67. A fluorescent sphere with a diameter of 10 μm was assumed as the object. The middle plane was imaged to evaluate the imaging property inside dense sample. The light sheet activation improves the contrast of the emission and the resultant image but brings about a trade-off between the axial resolution and field of activation. A scanned Bessel sheet with single-photon activation provides a large field of view, whereas the contrast improvement is not sufficient due to the sidelobes of the Bessel beam. Two-photon activation with a scanned Bessel sheet realizes the fluorescence distribution highly confined within the focal plane without sacrificing the field of view.

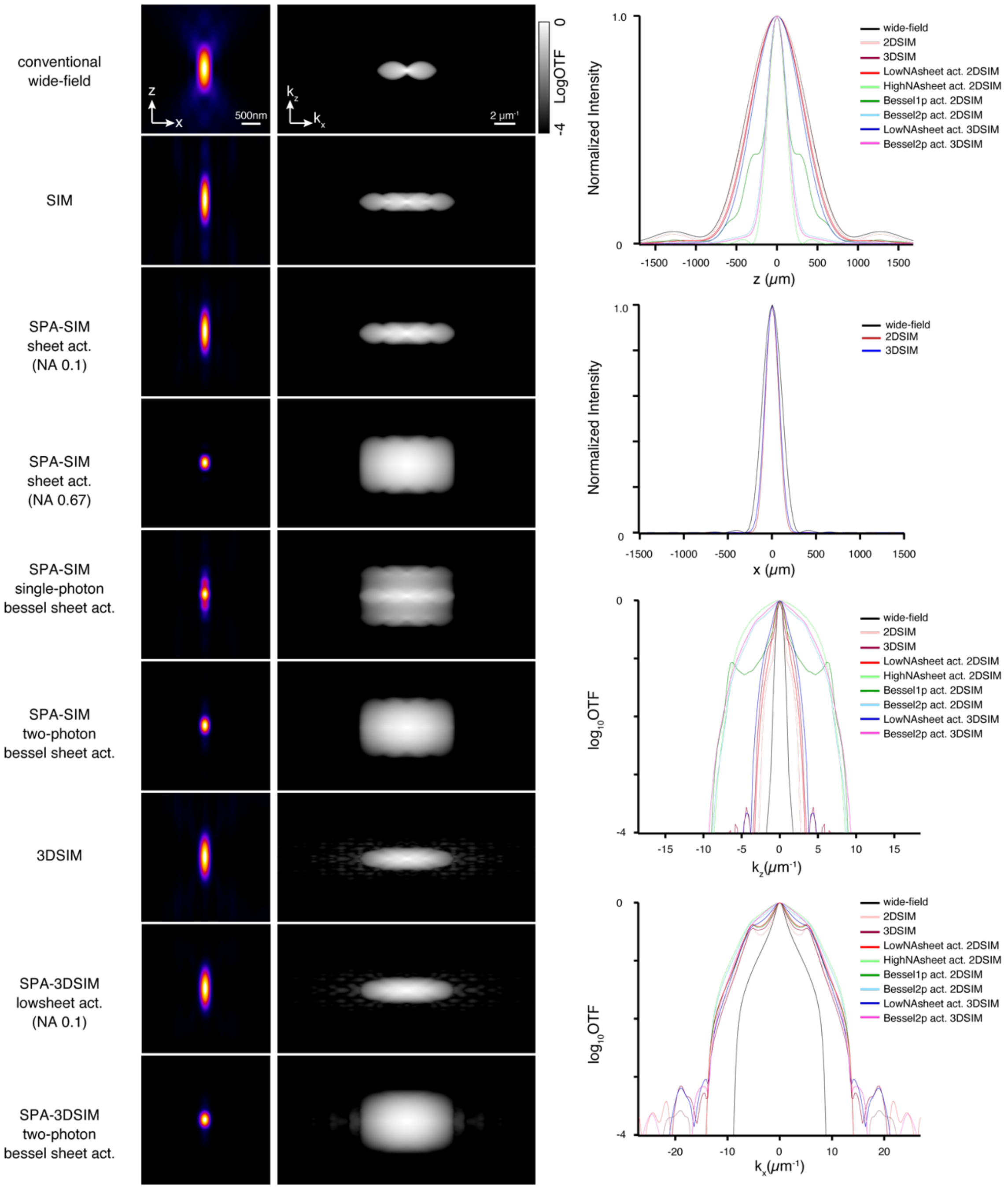


Fig. S3 Point spread functions and optical transfer functions of conventional wide-field microscopy, SIM, light sheet SPA-SIM with NAs of 0.1 and 0.67, Bessel light sheet for single- and two-photon activation in SPA-SIM with an NA of 0.67, 3DSIM, SPA-3DSIM with a light sheet and Bessel light sheet with two-photon activation. PSFs and OTFs were normalized by their individual maxima and plotted in linear and logarithmic scales, respectively. Intensity profiles at $x = 0$, $z = 0$, $k_x = 0$, and $k_z = 0$ were shown. We set the contrast of OTFs to a minimum value of -4 to eliminate calculation noise.

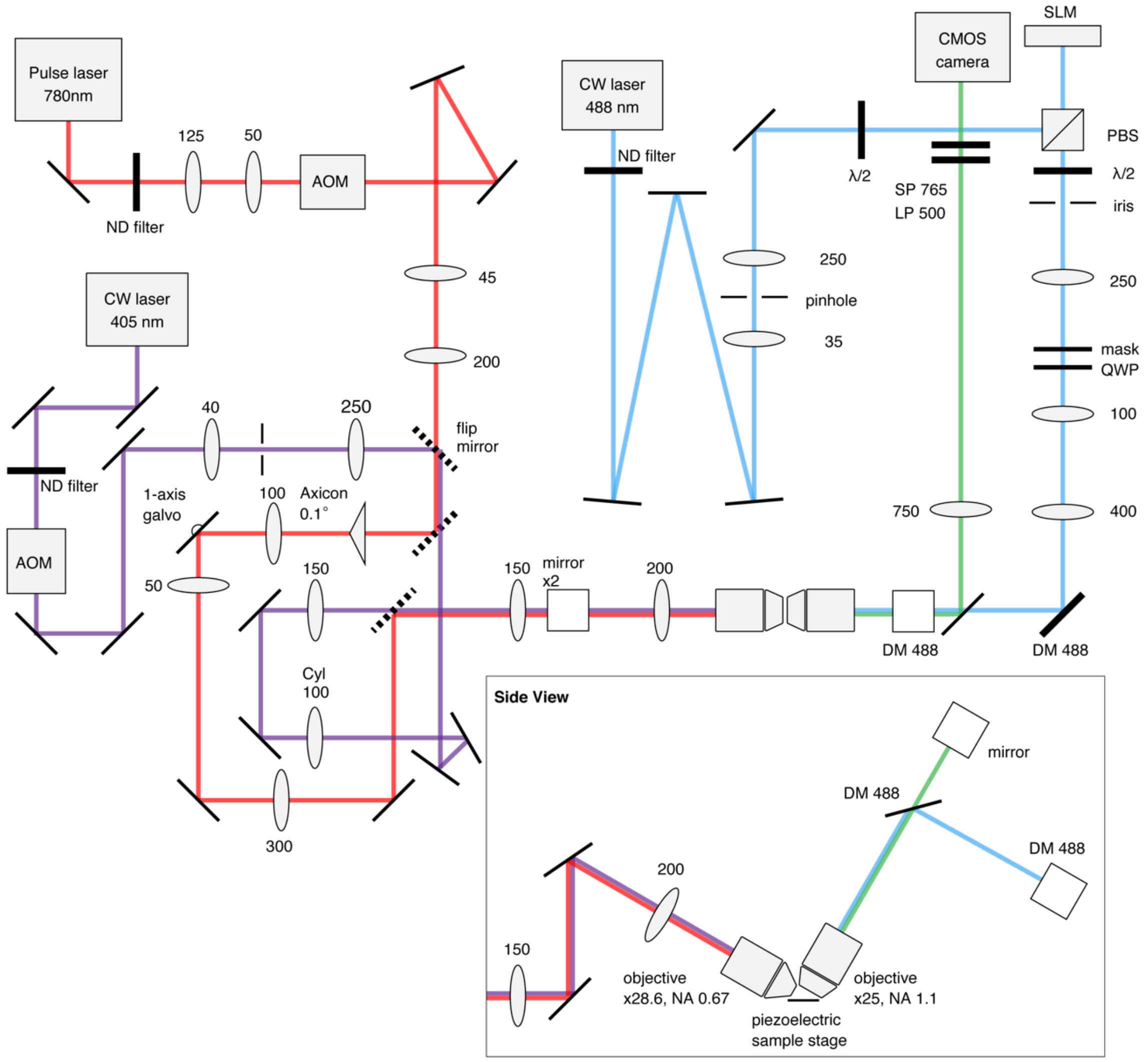


Fig. S4 optical setup

A continuous wave (CW) laser beam with a wavelength of 405 nm (Excelsior 405, Spectra Physics) or femtosecond pulse laser with a wavelength of 780 nm (MaiTai HP, Spectra Physics) is used for single- or two-photon activation. The pulse width and the repetition rate are 100 fs and 80 MHz, respectively. The laser beam is passed through an acousto-optic modulator (AOM, ASM-802B8, IntraAction Corp.) for digital modulation of the irradiation power to the samples and is collimated by two lenses with a spatial filter pinhole. The beam is shaped in two ways: a cylindrical lens or an axicon lens and a galvanometer mirror to form a light sheet on the sample plane. The laser beam was then focused using a water immersion objective lens (28.6×, NA0.67 model 54-10-7, Special Optics) to activate the sample. For excitation, a CW laser beam with a wavelength of 488 nm (Sapphire 488-500 LPX, Coherent) collimated by two lenses with a spatial filter pinhole is reflected by a polarized beam splitter (PBS, PBS251, Thorlabs) to a spatial light modulator (SLM, SXGA 3DM, Fourth Dimension Display) after passing through a λ/2 plate (WPH10M-488, Thorlabs). The SLM digitally modulates the beam pattern into sinusoidal patterns for structured illumination. After passing through PBS and a quarter-wave plate (QWP, WPQ-4880-4M, Sigma Koki) for circular polarization, the polarization conditions are adjusted for all structured illumination angles. 0th-order diffracted beam and other unwanted diffractions are blocked using a silver-coated mask in the 2DSIM mode. In the 3DSIM, a different patterned mask that can pass through a 0th-order beam is used. The beams passing through the mask are reflected by two long-pass dichroic mirrors (DM, Di03-R488-t3-25×36, Semrock) to compensate for the phase delay and were focused on the samples using a water objective lens for excitation (25×, NA1.1, CFI75, Nikon). The Fluorescence emission from the samples is collected using the same objective lens, passed through the DM, and imaged using a CMOS camera for detection (OLCA-Flash 4.0 v3, Hamamatsu Photonics). Long-pass (LP500, BLP01-488R-25, Semrock) and short-pass (SP765, FF01-790/SP-25, Semrock) filters are set in front of the camera to reject illumination and activation light. Nine or fifteen images are acquired with three angles and three or five phases of structured patterns for each super-resolution image. 1D piezoelectric stage (SFS-H40X, OptSigma) was used to move the samples for 3D imaging.

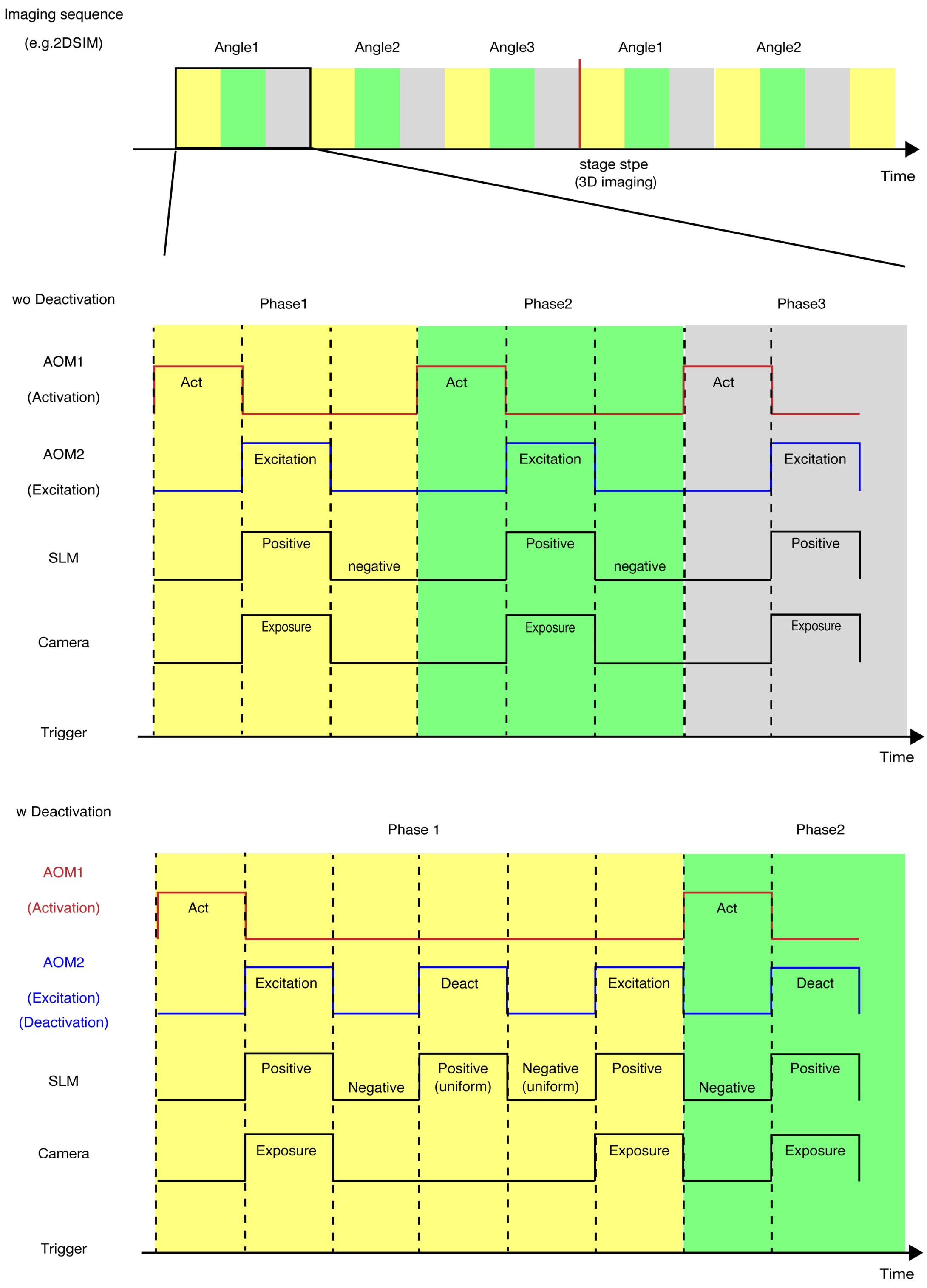


Fig. S5 Timing diagram of the measurement system. The top row shows the brief imaging procedure of SIM. Nine images with structured illumination of three angles by three phases are obtained to reconstruct super-resolution images. 3DSIM imaging requires five phases for each angle. For time-lapse imaging, the same process is repeated to fulfill the designated time slot. For 3D imaging, the sample stage moves a single step after a single SIM-imaging process. Detailed diagrams in cases of with and without deactivation are shown below. AOMs for excitation and activation, spatial light modulator, and camera are triggered by a PC, which is indicated by black dotted lines. The imaging process without deactivation, which is used in time-lapse or 3D imaging, consists of activation, excitation, and relaxation for SLM. Activation is separated from the excitation process to avoid the temporal cross-talk of fluorescence excited by activation light and excitation light. Excitation, SLM, and camera are triggered at the same time to obtain the fluorescence images under structured illumination. A negative pattern for structured illumination is applied to SLM for avoiding the burn-in. Deactivation with uniform illumination and another excitation step are added in another imaging mode to avoid the on-state proteins being left in the next measurement and observe the distribution of the fluorescence from off-state proteins.

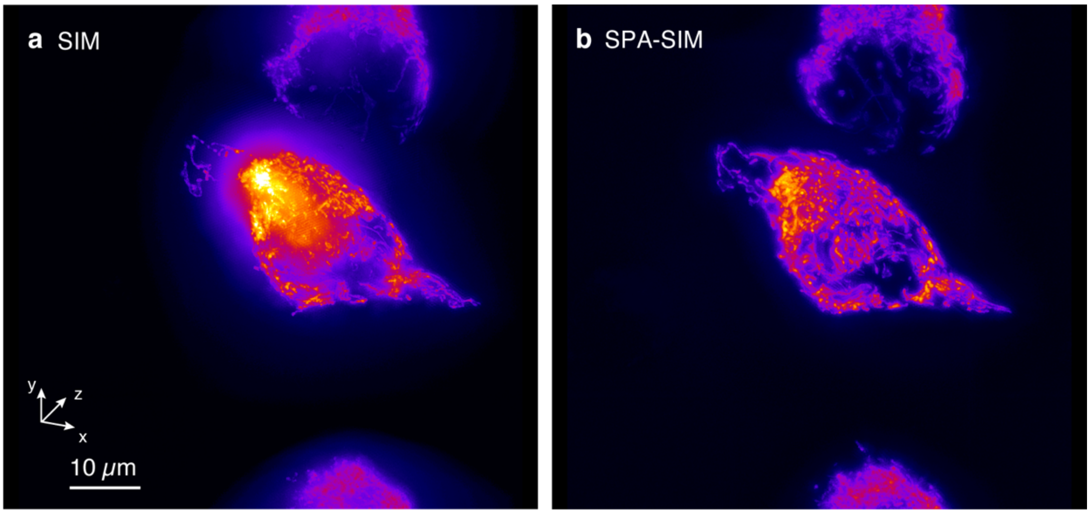


Fig. S6 3D images of mitochondria in living HeLa cells with (a) conventional SIM and (b) SPA-SIM with single-photon light sheet activation. The imaging volumes were 71 × 71 × 25 μm$^3$. The experimental conditions are shown in Extended Fig. 2. SPA-SIM was used to visualize the mitochondrial components distributed inside the cells owing to out-of-focus rejection.

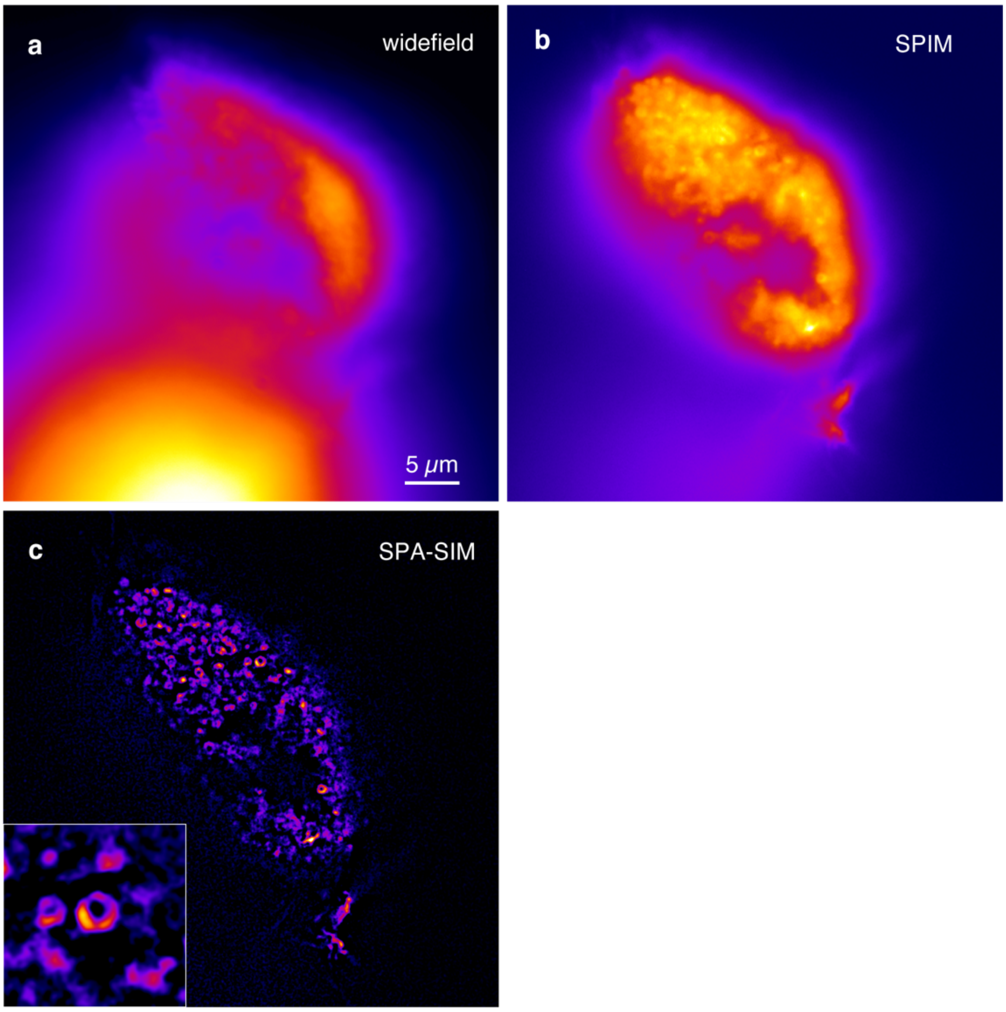


Fig. S7 Fluorescence images of the same MEF as Fig. 2k-l observed with (a) wide-field and (b) SPIM mode. (c) SPA-SIM image (Fig. 2k) is shown for comparison.

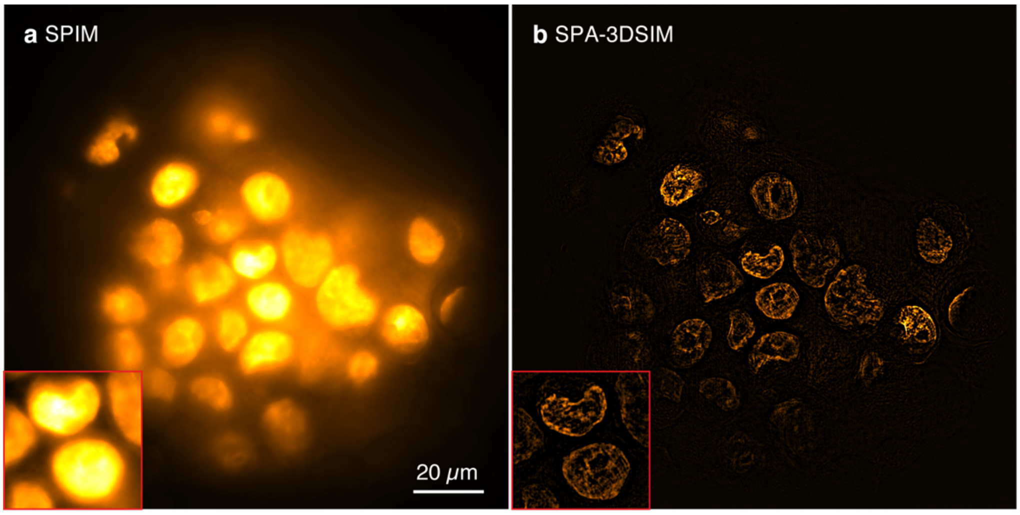


Fig. S8 Fluorescence images of the same cell spheroid sample as Fig. 2s-u observed with (a) SPIM. (b) SPA-3DSIM image (Fig. 2u) is shown for comparison. The reconstruction of the super-resolution image failed because the structured illumination pattern at the focus plane was not detected due to the high out-of-focus signal.

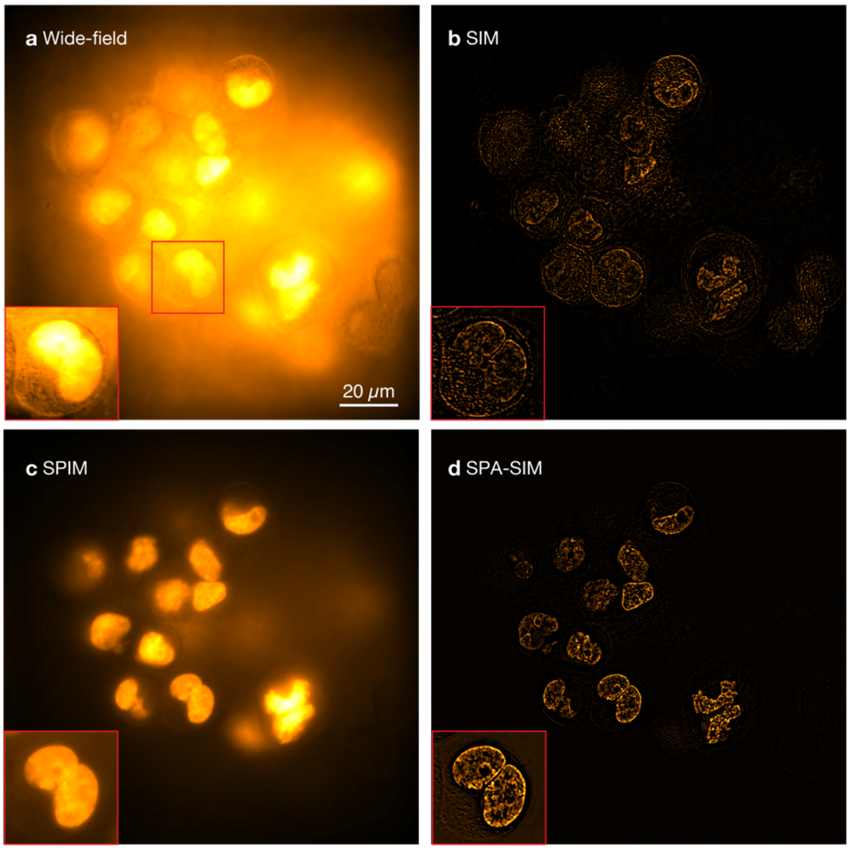


Fig. S9 Fluorescence images of nuclei labeled by rsGamillus-S in a cell spheroid with a diameter of ~100 μm at the depth of 25 μm observed with (a) wide-field, (b) SIM, (c) SPIM, and (d) SPA-3DSIM with two-photon activation by a Bessel sheet. Experimental conditions were the same as those shown in Fig. 2 (q)-(s).

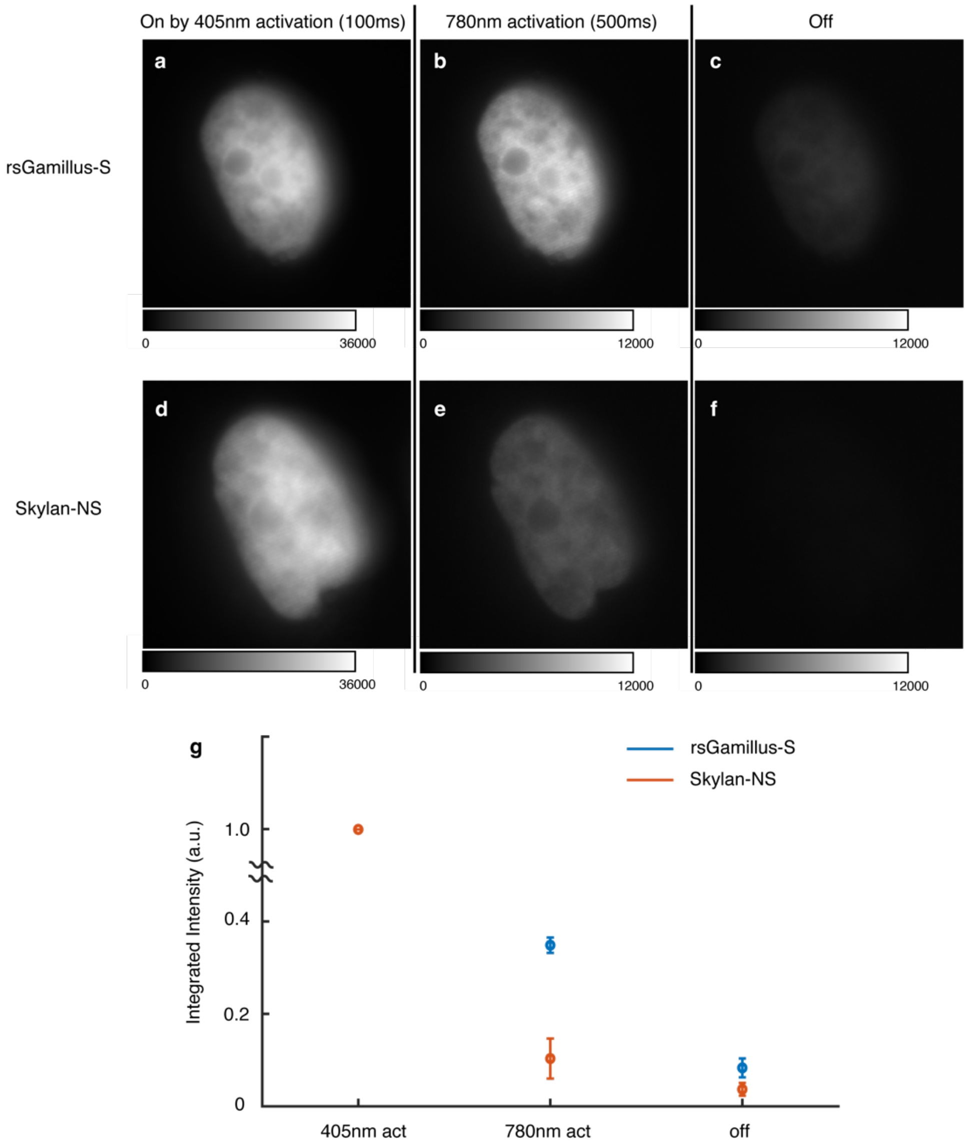


Fig. S10 Representative fluorescence images of histone H2B in a nucleus of a living HeLa cell labeled by rsGamillus-S and Skylan-NS. The images were captured after three different procedures as follows. (a,d) RSFPs were activated by 405 nm in 100 ms, switching almost all FPs to on-state. (b,e) Two-photon activation with Bessel sheet was applied in 500 ms which is same as the experimental conditions in the main text. (c.f) 488 nm light was irradiated for switching FPs to off-state. Activation laser (405 nm and 780 nm) powers were 248 μW and 67 mW, respectively, and 488 nm excitation with 6.05 (W/cm$^2$) was used for imaging. (f) Averaged integrated intensity of 5 fluorescence images of a cell labeled by rsGamillus-S and Skylan-NS. Different cells in the same dish were measured. Integrated intensities under different activation conditions were normalized by the value of 405 nm activation. We confirmed rsGamillus-S was 3.37 times more efficiently activated by two-photon Bessel sheet presumably due to the higher on-switching rate, which is suitable for SPA-SIM imaging.

Supplementary Video 1
Time-lapse images of actin filament movement in living a HeLa cell labeled with Skylan-NS observed by SPIM and SPA-SIM shown in Fig. 2 (g). The acquisition rate was 0.6 frame/s with an interval of 7 s to visualize slow actin motion.

Supplementary Video 2
Time-lapse images of mitochondria in a living HeLa cell labeled with Skylan-NS observed by SPIM and SPA-SIM at 1 fps with an interval of 4 s. Several images at different time points and enlarged views are shown in Fig. 2 (h).

Supplementary Video 3
3D images of mitochondria in a living HeLa cell labeled with Skylan-NS obtained using 3DSIM and SPA-SIM. The images from a single viewpoint are shown in Fig. 2 (i,j). The imaging volumes were $71 \times 71 \times 25\ \mu m^3$.

Supplementary Video 4
3D images of mitochondria in living HeLa cells labeled with Skylan-NS obtained using conventional SIM and SPA-SIM. The images from a single viewpoint are shown in Fig. S8. The imaging volumes were $71 \times 71 \times 25\ \mu m^3$.
